# Genome-wide identification and characterization of DNA/RNA differences associated with *Fusarium graminearum* infection in wheat

**DOI:** 10.1101/2021.06.25.449684

**Authors:** Guang Yang, Yan Pan, Ruoyu Zhang, Jiaqian Huang, Wenqiu Pan, Licao Cui, Weining Song, Xiaojun Nie

## Abstract

RNA editing (DNA/RNA differences) as a post-transcriptional modification approach to enrich genetic information, plays the crucial role in regulating diverse biological processes in eukaryotes. Although it has been extensively studied in plant chloroplast and mitochondria genome, RNA editing in plant nuclear genome, especially those associated with Fusarium head blight (FHB), is not well studied at present. Here, we investigated the DNA/RNA differences associated with FHB through a novel method by comparing the RNA-seq data from *Fusarium*-infected and control samples from 4 wheat genotypes. A total of 187 DNA/RNA differences were identified in 36 wheat genes, representing the first landscape of the FHB-responsive RNA editome in wheat. Furthermore, all of these 36 edited genes were located in the FHB related co-expression gene modules, which may involve in regulating FHB response. Finally, the effects of DNA/RNA differences were systematically investigated to show that they could cause the change of RNA structure and protein structure in edited genes. In particular, the G to C editing (chr3A_487854715) in TraesCS3A02G263900, which is the orthology of *OsRACK1*, resulted that it was targeted by *tae-miR9664-3p* to control its expression in different genotype through different editing efficiency, suggesting RNA editing could mediate miRNA to participate in the regulation network of FHB tolerance. This study reported the first wheat DNA/RNA differences associated with FHB, which not only contribute to better understand the molecular basis underlying FHB tolerance, but also shed light on improving FHB tolerance through epigenetic method in wheat and beyond.

## Introduction

Wheat is considered as one of the most important staple crops all over the world, which accounts for approximately 30% of the global cultivated area, and provides 20% of the world’s food consumption (Shewry 2009). Wheat is also an important source of human protein and mineral elements intake (Gill et al. 2004; Appels et al. 2018). Continuous increased and stable production of wheat holds the promise for ensuring global food security under the challenge of population booming and limited resource input in future (Miransari and Smith 2019). Fusarium head blight (FHB), that is also called scab and caused mainly by *Fusarium graminearum*, is one of the most destructive diseases of wheat, resulting in huge loss of wheat yield and also imposing great health threats on both human beings and livestock due to the DON toxin (Bai and Shaner 1994; Dexter et al. 1996). More seriously, fusarium head blight has gradually become the major hazard and limitation of wheat production in recent years because of the climate change and the expansion of conservation agriculture (Zhu et al. 2019). Thus, revealing the mechanism of FHB resistance and then breeding for FHB-tolerant wheat varieties is crucial to cope with these problems. Extensive studies have been carried out to survey resistant germplasm, map and locate the QTLs, together with clone the major functional genes as well as illuminate the regulation mechanisms of FHB response in wheat (Buerstmayr et al. 2009; Rawat et al. 2016; Jia et al. 2018). The great breakthrough is the cloning and functional validation of the *Fhb1* (syn Qfhs.ndsu-3BS) from cv. Sumai No.3, which is widely used in breeding practice (Li et al. 2019; Su et al. 2019), as well as the *Fhb7*, which was horizontally transferred from fungus in wheat (Wang et al. 2020). Additionally, based on RNA-seq technology, the gene expression profiles and gene co-expression network analysis have also been systematically performed to identify the FHB-responsive genes and to discover regulators and genes associated with constitutive resistance (Pan et al. 2018; Hofstad et al. 2016).

RNA editing (DNA/RNA difference) is a conserved post-transcriptional modification mechanism that base change or modification is occurred when DNA transcribed into RNA molecule (Keller et al. 1999; Stern et al. 2010). Together with alternative splicing (AS), RNA editing process provides the indispensable approach for enriching the genetic information and diversifying the transcriptome, which plays the vital role in growth and development as well as stress tolerance in many organisms (Wang et al. 2016). Previous studies found that up to 55% of the genetic information in the mature mRNA molecules were inconsistent with the initial DNA sequence (Takenaka et al. 2013; Wakasugi et al. 1996). RNA editing was firstly identified in the mitochondrial genome of trypanosome in 1986, and now it has been widely reported in many species, including animals, plants as well as fungi (Bock et al. 1994; Drescher et al. 2002; Liu et al. 2016). In mammals, the common type of RNA editing is the deamination of adenosine (A) to inosine (I), which is mainly mediated by the specific ADAR (adenosine deaminase acting on RNA) family of enzymes (Savva et al. 2012). At the same time, A to I conversion, independent of ADAR enzyme, is also identified in fungi. In plants, which is lacking the ADAR gene family, RNA editing was mainly found in the organelle genome through bioinformatic prediction and molecular cloning approach, and they were generally regulated by pentapeptide repeat (PPR) domain protein family (Drescher et al. 2002; Shikanai 2006). With the advances in high-throughput sequencing, RNA-seq technology provides an efficient, unbiased and economic way to identify RNA editing on a genome-wide scale. Using this method, a large number of studies have been conducted to study the RNA editome or landscape in human and other model species, illuminating the prevalence and importance of RNA editing (Peng et al. 2012). However, the study of RNA editing in plants is lagging behind, especially genome-wide identification of DNA/RNA differences in plant nuclear genome only performed in Arabidopsis up to now (Meng et al. 2010). The plant-pathogen system provides an ideal model to identify RNA editing targets associated with pathogen based on RNA-seq method, in that the transcriptome sequences of the pathogen-treated samples and the counterpart control samples of the same genotype are generally produced, so as to exclude genotype-specific polymorphisms and mutations to ensure the accuracy of DNA/RNA difference identification.

Here, we systematically investigated the DNA/RNA differences of wheat in response to *F. graminearum* using the publicly available RNA-seq samples of four wheat genotypes (Nyubai, Wuhan 1, HC374, and Shaw), at 2 and 4 days post inoculation (dpi) with *F. graminearum* infection to understand the roles of DNA/RNA differences in regulating FHB tolerance in wheat. This study not only identified the DNA/RNA difference sites associated with FHB resistance to enrich the epigenetic mechanism of FHB response in wheat, but also pave the way to investigate RNA editiome using RNA-seq in wheat and beyond.

## Materials and methods

### RNA-seq data and Reads Mapping

The transcriptional dynamics associated with resistance and susceptibility against FHB of four wheat genotypes were performed by Pan et al (Pan et al. 2018). A total of 48 RNA-seq data of wheat spikes were provided by this study and publically available from the Sequence Read Archive (SRA) database with the accession no. of SRP139946. These datasets were downloaded and used in this study, including four wheat genotypes inoculated with water and *Fusarium graminearum* (strain DAOM233423) with 3 biological replicates at 2dpi and 4dpi for each genotype, respectively. Then, the raw RNA-seq reads were filtered for contamination with adaptor reads, low-quality reads, or unknown nucleotides using FastQC (version 0.11.8) and Trimmomatic (version 0.39). The cleaned RNA-seq reads were mapped against the reference genome (IWGSC RefSeq version 1.1) (Appels et al. 2018) using 2-pass mode of STAR (version 2.7.5c) (Dobin et al. 2012). The alignments were used for transcript assembly with StringTie (version 1.3.5). Furthermore, we quantified the read coverage of each gene by HTSeq (version 0.11.2) (*htseq-count -f bam -m union ${f}_sorted.bam $ref > ${f}count.txt*). Differentially expressed genes were identified using DESeq2 tool with the adjusted P value was less than 0.05 and |log_2_FoldChange| > 0 (Love et al. 2014).

### DNA/RNA difference sites identification

Firstly, using the MarkDuplicates tool of Picard (http://picard.sourceforge.net/) marked the repeated sequence in the bam files obtained by STAR (2-pass mode). Then, the reads on the exon were separated by using the SplitNCigarReads tool in GATK, and the N error base was removed and the read in the intron region was removed. HaplotypeCaller tool in GATK (Genome Analysis Toolkit) software was used to call SNPs with the parameter as follow: --*genotype_likelihoods_model ‘SNP’, --stand_call_conf ‘30’, --stand_emit_conf ‘30’* (Ramaswami et al. 2013).Then, the SNP was obtained as the raw gVCF file of each sample, and further used for subsequent analysis. To obtaining high confidence sites, we filtered raw VCF files step by step as follow: (1) Systematic error of the sequencing platform and software were corrected by GATK VariantFiltration tool, and we select the initial filter parameter *-filter “FS > 30.0”, -filter “ QD < 2.0*”; (2) To improve the accuracy, the three biological replicates were intersected to obtain DNA/RNA differences that appeared in three replicates simultaneously and the each sequence information was verified by Integrative Genomics Viewer (IGV); (3) To avoid genotype-specific genomic SNP polymorphisms, we compared the DNA/RNA differences between *Fg*-treated samples and their counterpart control samples of the same genotype, and the same DNA/RNA differences between them were removed.

Finally, the 654,653 SNP variations of 1,002 wheat genotypes include 717 genotyped by DARTseq platform and 285 genotyped by Wheat 660K SNP array (Zhou et al. 2018) were used to map the qualified DNA/RNA differences obtained from above analysis to filter out the putative SNP sites with the same genomic physical position. Through these programs, the accuracy and high-reliability DNA/RNA differences were finally obtained. They were annotated with SnpEff tool (version 3.6) with the annotation file downloaded from Ensemble Plants database (http://plants.ensem-bl.org/index.html). The orthologous genes of candidates in *Arabidopsis* or rice were also obtained from Ensemble database.

### Co-expression network analysis

Gene co-expression network analysis was conducted based on the all genes using the R package WGCNA tool (Langfelder and Horvath 2008). Genes with an average TPM value greater than 2 and at least one sample expressed were used. A Pearson correlation coefficient matrix was computed. Then, we calculate log_10_[p(k)] and log_10_(k) separately, and fit the calculated results to determine 6 is easier to meet the criterion of negative correlation between log_10_[p(k)] and log_10_(k). After determining the beta value, we converted the relationship matrix into an adjacency matrix and TOM similarity matrix was generated for each adjacency matrix. The different coefficients and hierarchical clustering trees of different nodes are calculated and constructed; hierarchical clustering was employed based on the similarity matrix to cluster genes. To obtain the correct module number and clarify gene interaction, we set the restricted minimum gene number to 30 for each module and used a threshold of 0.25 to merge the similar modules. Genes that have higher weight in important modules were chosen to constructed co-expression network. The traits data publicly available, including *Fg* treatment, *Fg* time, *Fg* percent, *Fg* GAPDH (Glyceraldehyde-3-phosphate dehydrogenase) and DON (Deoxynivalenol) were used for trait-module correlation analysis. GO and KEGG enrichment analysis was conducted using KOBAS 3.0 software (Xie et al. 2011) with the annotation file of *Arabidopsis thaliana* as background.

### RNA structure analysis

RNAfold in the Vienna RNA Secondary Structure Package (Gruber et al. 2008) were used to predict the secondary structure of candidate RNA editing genes before and after editing. In order to compare the RNA structure of different genes reasonably, we calculated the normalized free energy of RNA secondary structure by the method of predecessors (Mao et al. 2013). Each candidate sequence was randomly shuffled 100 times to control base composition before and after editing. Then, normalized minimum folding free energy (MFE) of each candidate was calculated using RNAfold by

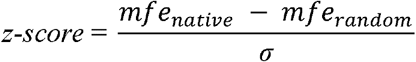

Among equation, *mfe_native_*, *mfe_random_* and σ is the free energy of native sequence, mean MFE of 100 random sequences and standard deviation of the MFE of 100 random sequences, respectively.

### miRNA target analysis

To determine whether DNA/RNA differences affected miRNA targeting sites, all of candidate editing genes transcripts were searched against the publish wheat miRNAs in the miRBase using psRNATarget tools (Dai and Zhao 2011) to predict whether they targeted by miRNAs. The possibility of miRNA targeted on edited genes was scored using Schema V2 (2017 release) schema, and selected the result with the minimum expected value as the optimal prediction.

### Protein domain and structure analysis

The PFAM database (33.0 release) were used to predicted protein domain by HMMER v3.3.1 tools (Finn et al. 2011) with E-value < 1×10^−5^. Protein 3D Structure was predicted using homology modeling methods in SWISS-MODEL database (https://swissmodel.expasy.org/). We selected the best result that the model has the highest agreement with the target protein and is greater than 30%.

### Orthologous gene analysis

In order to better clarify the potential function of the RNA editing genes, the functionally validated genes in Arabidopsis and rice were downloaded from the TAIR (https://www.arabidopsis.org/) and Ricedata (https://www.ricedata.cn/gene/) databases respectively. Then, we used validated genes as the query to search against the local wheat proteins by BLASTP tool (Camacho et al., 2009) with the identity more than 40% and E-value of 1e-10 as threshold.

## Results and Discussion

### Identification of DNA/RNA difference using RNA-seq data

Based on the RNA-seq data, a total of 137,037 transcripts and 110,777 gene loci were constructed for the four wheat genotypes, which covered more than 99% of the reference genome of wheat IWGSC V1.1 (Fig. 1a). Then, the difference of sequence between RNA and DNA were identified using the method described above, and 16,399 putative DNA/RNA difference sites associated with FHB were obtained, suggesting massive difference events were occurred in wheat responding to FHB. In detail, a total of 292 difference sites in wheat genotype HC374, 444 in Nyubai and 3,490 in Wuhan 1 as well as 3,836 in Shaw were found at 2 dpi, while at 4dpi 568, 490, 3,847 and 3,432 sites were identified in HC374, Nyubai, Wuhan 1 and Shaw, respectively (Fig. 1b) (Table S1). Compared to 2dpi, the DNA/RNA difference sites at 4dpi always showed more abundant in all of three resistant genotypes, indicating that the number of difference site increased with the extension of *Fg* injection in resistant genotypes while no found in susceptible genotype Shaw. Furthermore, 14,452 unique DNA/RNA difference sites presenting in 8,346 genes were obtained through removing the redundant sites, of which 5,361 genes have one sites, follow by 1,801, 633 and 244 genes with 2, 3 and 4 sites, as well as 307 genes with more than 5 sites. Further studies to decipher the molecular basis underling DNA/RNA difference could provide vital clues for the complex of transcription regulation as well as genetic variations. The physical position of these difference sites mainly located about 10kb upstream or downstream of the TSS of the corresponding correlation genes (Fig. 1c), suggesting that difference events may have a large influence on these genes’ expression. Finally, we further annotated these DNA/RNA difference sites. Results showed that 9,306, 206, 900 and 1,926 sites were located in CDS, intron, 3’UTR and 5’UTR, respectively. A total of 6,586 and 2,720 DNA/RNA difference sites emerged as missense variant and synonymous variants, accounting for 45.57% and 18.82% respectively, suggesting that missense variants would lead to one amino acid change in the protein composition (Table S1). At the same time, there also 426 sites could cause the amino acid substitution, particularly, 388 stop-gained variations were identified. Although the DNA/RNA difference associated with FHB has been identified, the false positive results still appeared because of the coverage of reads and the existence of SNP. Therefore, further validation analysis was needed to obtain reliable RNA editing sites related to FHB from DNA/RNA differences.

### Identification of putative DNA/RNA differences associated with FHB tolerance

Based on all DNA/RNA differences results, we conducted a comprehensive screening of FHB-related RNA editing sites using the IGV tools, and we focused on the common RNA editing sites of four varieties and three resistant varieties to eliminate false positives caused by genotype differences (see **Materials and methods** for more details). In detail, a total of 206 RNA editing sites were identified in two stages and four wheat genotypes, of which contained 187 unique RNA editing sites in 36 genes (Fig. 1d and Table S2). Among them, 159 sites were common in four wheat genotypes and 47 were common in three resistant genotypes. Compared to 2dpi, the editing sites at 4dpi always showed more abundant, indicating that the number of RNA editing sites increased with the extension of *Fg* injection in each genotype. Moreover, 13 common RNA editing events of TraesCS2D02G179300, TraesCS2D02G405500, TraesCS3A02G263900, TraesCS4A02G107600 and TraesCS5A02G073800 were found in 2dpi and 4dpi in four wheat genotypes, and 4 common RNA editing sites of TraesCS5A02G073800 were identified in 2dpi and 4dpi in three resistant genotypes (Table S2). These loci may play an important role in different stages of FHB response. From the perspective of editing type (Fig. 1e), 95 editing sites (50.79%) were the type of transition, of which the conversion between C and T accounted for 24.06%, and A and G accounted for 26.74%, respectively, representing the two most abundant editing types. These two types were also the two canonical RNA editing (Pachter 2012). Among transversions, G to T (10.70%) and C to A (9.63%) were the most abundant, following by C to G, G to C, T to G and A to C with all of the value of about 23.53%, while T to A and A to T types are the lowest ones with the value of 2.67%, respectively. It is obvious that base transition events were significantly lower than transversion in these editing sites (Transition/Transversion ratio was 1.033) although there are twice as many possible transversions on the fact of frequency. It is well known to us that transitions are enriched over transversions at genome level as transversions generally result in the amino acid substitution and are more likely to be depleted due to evolutionary selection (Guo et al 2017). Then, we further annotated these RNA editing sites. Results showed that 162 and 25 sites were located in protein coding region and none coding region, respectively. A total of 45 and 117 editing sites emerged as missense variant and synonymous variants, accounting for 24.06% and 62.57%, respectively (Fig. 1f), of which the editing sites could cause the amino acid substitution might have important regulation roles in response to *Fg* infection in wheat. Editing efficiency was reflected by the ratio of edited reads to total reads of each edited sites. The RNA editing efficiency of each variety was significantly different between control group and treatment group, of which the editing efficiency of Shaw was the highest (Fig. 2a). The density of efficiency of three resistant varieties showed left skewed distribution and Shaw showed right skewed distribution (Fig. 2b). The difference of editing efficiency between different varieties may indicate the difference of FHB tolerance.

**Fig. 1.**
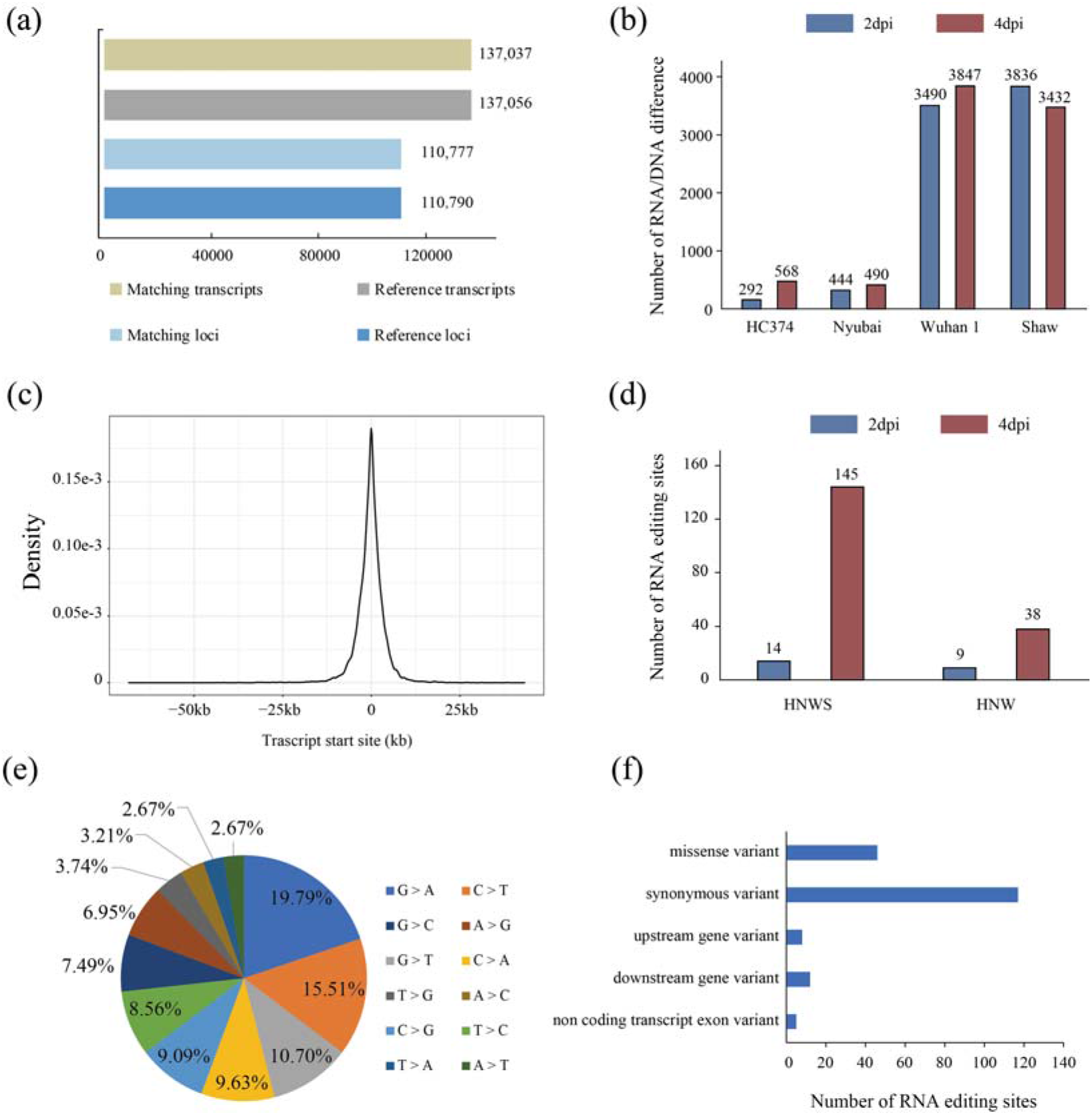
Characterization of DNA/RNA difference sites and RNA editing sites. (a) The assembled genes and transcripts based on the RNA-seq using in this study against the wheat reference genome IWGSC v1.1. (b) The numbers of DNA/RNA difference sites were identified in four wheat varieties at 2dpi and 4dpi, respectively. (c) The distribution of DNA/RNA difference sites distance from TSS (transcription start site) of its related genes. (d) The number of RNA editing sites shared by four varieties (HNWS: HC374, Nyubai, Wuhan 1, Shaw) and three resistant varieties (HNW: HC374, Nyubai, Wuhan 1). (e) Distribution of all unique RNA editing site types. (f) Distribution of RNA editing sites by transcription regions. The y axis represents the different types of regions, and the x axis shows the abundances of RNA editing sites

**Fig. 2.**
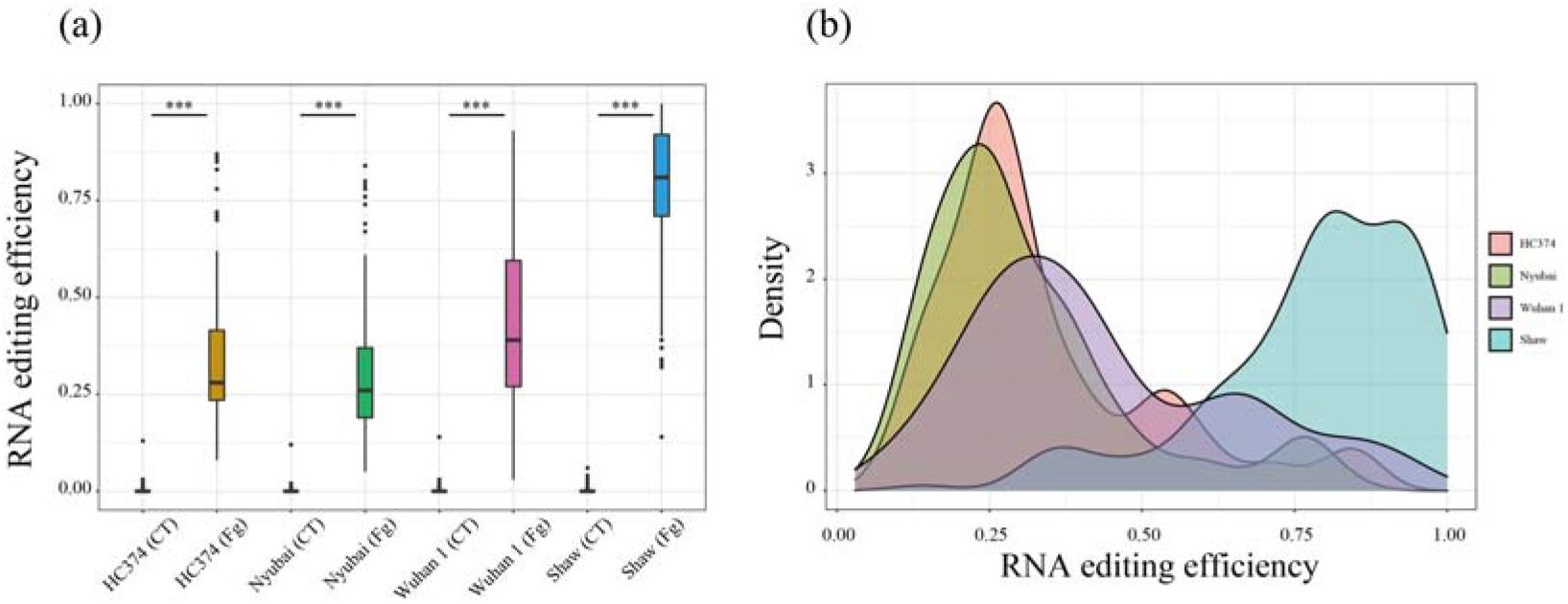
Comparison of RNA editing efficiency in different varieties (a) and the distribution of RNA editing efficiency (b).

### Integration of RNA editing sites and gene expression

To further confirm the FHB-responsive RNA editing sites, we investigated the expression patterns of these edited genes (Fig. 3) (Table S3). Among them, TraesCS1A02G258800 and TraesCS1D02G258800 was differential expressed in each stage of four varieties. Meanwhile, four differential expressed edited genes (DEEGs) were shared by all of the four varieties in 4dpi. Compared with sensitive variety Shaw, TraesCS3D02G328300 showed differential expression in 2dpi of three resistant varieties, indicating the potential function of this gene in response to FHB. On the contrary, six and three genes of Shaw were down regulated and up-regulated respectively, and there was no difference in the expression of these genes in resistant varieties after inoculation. Furthermore, TraesCS4D02G319400 is annotated to encode a glycosyldehyde-3-phosphate dehydrogenase (GAPDH). It has been demonstrated that GAPDH involved in the protein aggregation and DNA repair due to stress-related factors (Zaffagnini et al. 2019), indicating that RNA editing in GAPDH might mediate glycolysis pathway to promote the FHB tolerance. Otherwise, the differential expression of TraesCS3A02G263900 was found in 4dpi of Shaw, indicating the potential function associated with *Fg* infection. Meanwhile, *OsRACK1* was the orthologues of TraesCS3A02G263900 and it has been proved to have the function of resistance to rice blast (Nakashima et al. 2008).

**Fig. 3.**
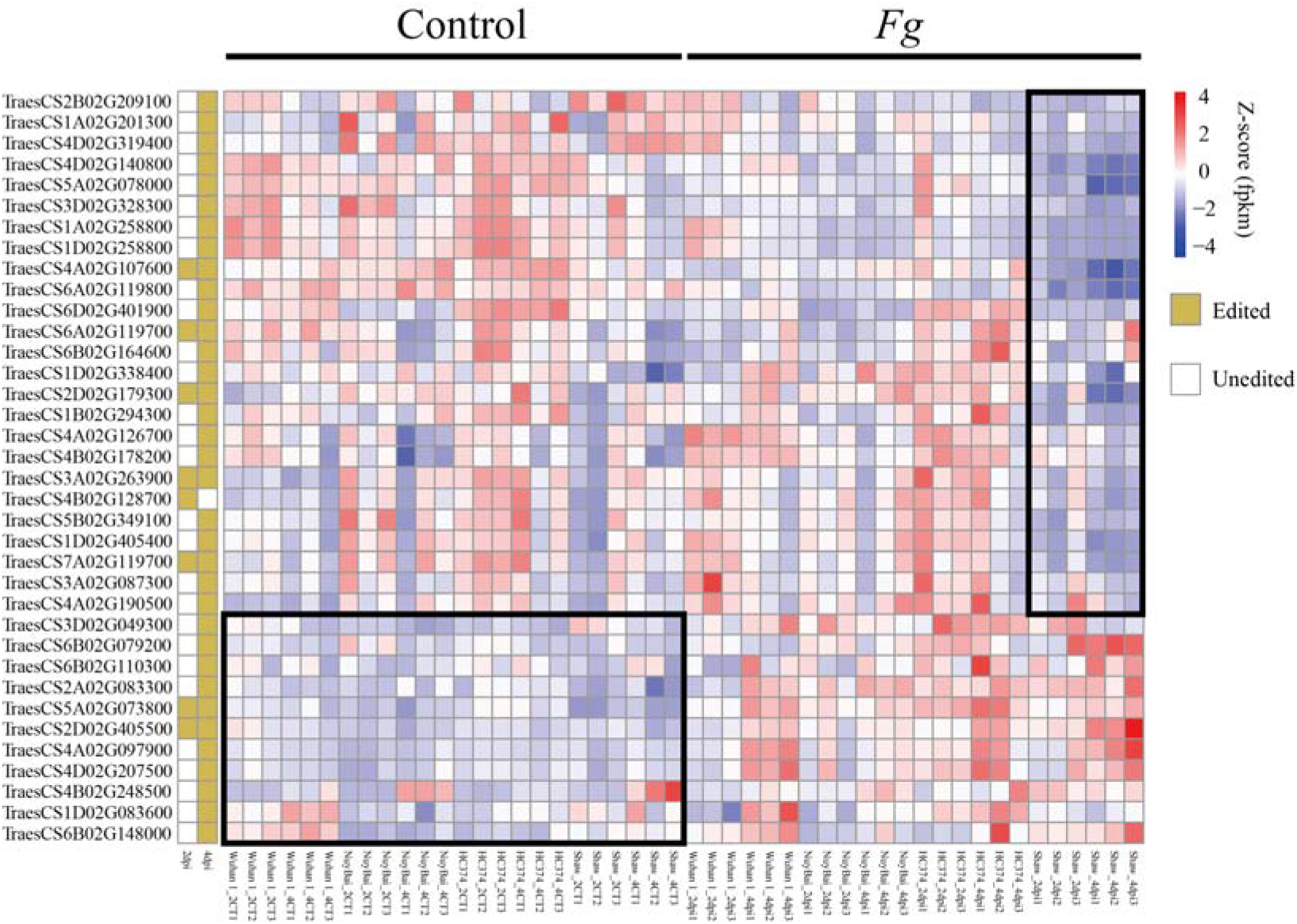
The expression profiles of the 36 candidate RNA editing genes among resistant and susceptible genotypes at two inoculation time points. The left block diagram represents whether RNA editing occurred in these genes. Blank means no editing, while green means editing. The middle heatmap represents the expressions of these genes in four varieties. CT: Control group. dpi: days post inoculation

To preliminarily understand the function and regulatory network of these RNA editing genes, we further constructed the WGCNA co-expression network based the 58,280 expressed genes and then linked the co-expression modules with the available phenotypic data of the *Fg* infection, including percentage (*Fg* infection), DON (Deoxynivalenol), GAPDH (Glyceraldehyde-3-phosphate dehydrogenase RNA level) and infection time, which were referred from previous study (Pan et al. 2018). Totally, 34 co-expression gene modules were obtained by constructing a scale-free network and dynamic tree cutting (Hierarchical Cluster) and modules were renamed M1-M34 according to the number of module genes, of which M1 module contained 12,301 genes, ranking the largest module, followed by M2 with the 7,217 genes, while M34 modules had only 39 genes (Fig. S2). Furthermore, Pearson correlation coefficient matrix was calculated between the modules and phenotypes (Fig. S3 and Table S4). Results showed that the M2 and M11 had high positive correlations with all the 5 phenotypes about *Fg* infection, which might be the key module associated with *Fg* infection. And M12 and M33 had positive correlations with four phenotypes (percentage, DON, GAPDH, infection time). M5, M16, M29 and M32 had positive correlations with *Fg* infection and infection time.

Then, we identified the co-expression module of each editing gene (Table S4). M1 module contained most of the editing genes, following with M4. The co-expression modules containing editing genes were associated with at least one FHB responsive trait. It is worth noting that five co-expression modules (M1, M2, M5, M7, M11) of the candidate genes were positively or negatively correlated with Fg infection, indicating that these genes play a more important role in wheat scab response. GO enrichment analysis of the edited genes found that most genes (77.78%, 28) were enriched in the term of cytosol (GO:0005829, 4.36E-23), and 5 genes enriched in defense response to fungus (GO:0050832, 2.20E-06) (Table S5). At the same time, the candidate genes were also enriched into the terms related to structure of protein or RNA, such as mRNA binding (GO:0003729, 2.82E-21), cellular response to unfolded protein (GO:0034620, 2.69E-11), misfolded protein binding (GO:0051787, 5.41E-10), cell wall (GO:0005618, 1.59E-06) (Table S5). For KEGG pathway enrichment, 3 genes (TraesCS4B02G178200, TraesCS3D02G328300, TraesCS4A02G126700) were found to enriched in MAPK signaling pathway-plant (ath04016, 4.25E-04), which is a crucial pathway related to abiotic and biotic stress (Zhang and Klessig 2001; Meng and Zhang 2013; Pitzschke et al. 2009). In addition, these 3 genes were also significantly enriched in plant-pathogen interaction (ath04626, 8.37E-04). These results suggested RNA editing was widely occurred in the genes associated with *Fg* infection, responding and tolerance in wheat. Further functional study of these RNA editing sites will not only mine some vital resistance gene for genetic improvement, and also contribute to enrich the epigenetic mechanism of FHB response in wheat.

### The effect of RNA editing on RNA structure

RNA structure is crucial to its function that RNA mainly depends on its local structure to interact with other proteins or molecules (Wan et al. 2011; Dethoff et al. 2012). The secondary structure of mRNA is mainly involved in cell processes through two forms: specific secondary structure binding to other molecules and conserved structural protective functional elements (Keller et al. 2012). RNA editing events directly affect the secondary structure of RNA (Solomon et al. 2017). Therefore, the RNA editing events in response to FHB may lead to changes in RNA structure and affect its function. Thus, RNA secondary structures of these FHB-responsive edited genes were predicted by the minimum free energy model. Results showed that 162 candidate sites in 32 editing genes could result in the change of RNA secondary structure (Table S6). After editing, the average MFE value were basically the same as that of before editing, but the average normalized MFE values had differences. Among these sites, the minimum free energy of 75 sites increased after editing, while that of the other 87 sites decreased. The normalized MFE of chr6B_55695035 site in TraesCS6B02G079200 increased by 40%, ranking the highest change. Meanwhile, TraesCS6D02G401900 had the minimum normalized MFE after chr6D_470684693 site editing. In the minimum free energy model, organisms will fold RNA into a secondary structure with minimum free energy, thus saving energy (DAWSON and YAMAMOTO 1999; Mathews et al. 1999). Therefore, MFE can be used to measure the stability of structures that the structure with low MFE value showed more stable. According to our prediction, 75 candidate RNA editing sites could lead to the instability of RNA, and then impair their normal function. On the contrary, the other 87 editing sites could lead the decrease of MFE value of the responding RNA secondary structure, indicating these editing sites played the crucial roles in maintaining or increasing the stability of their structure to perform their functions. These results suggested that RNA editing could impact on the function of the target genes through regulating their secondary structures. Further study the specific roles of RNA editing playing in regulating RNA secondary structure when in response to *Fg* infection might contribute to the genetic basis underling FHB tolerance.

### The effect of RNA editing on binding ability and protein structure

It has been demonstrated that RNA editing as the conserved post-transcriptional modification mechanism, could impact on binding ability, protein composition and protein structure (Takenaka et al. 2013). microRNAs (miRNAs) are one class of non-coding RNA to regulate gene expression through mediating targeted mRNAs cleavage or translational inhibition (Meng et al. 2010). RNA editing generally caused the mRNA sequence variations, which could impact on miRNA-mRNA binding (Mao et al. 2018). To better understand the function of RNA editing under *Fg* infection, we further investigated its effect on miRNA targeting. We identified seven RNA editing sites (1: chr3A_487854715; 2: chr3A_487854745; 3: chr3A_487854754; 4: chr3A_487854757; 5: chr3A_487854758; 6: chr3A_487854760; 7: chr3A_487854763) occurring in TraesCS3A02G263900 (Fig. 4a and Fig. S4), of which six sites were common to four varieties at 4dpi and site 2 were shared by four varieties at 2dpi and 4dpi (Table S2). Meanwhile, site 1 changed the amino acid from Glu to Asp and site 4 and 5 changed the common amino acid from Ala to Gly. Through analyzing the binding ability of gene after editing, we found that the occurrence of site 1 editing made the gene having the binding site of *tae-miR9664-3p* (Fig. 4a and Table S7) and the RNA secondary structure of gene was also changed by this RNA editing sites, of which the MFE of structure changed from −389.70 to −385.70 kcal/mol (Fig. 4b and Fig. 4c), indicating the stability of RNA structure was decreased. Interestingly, the editing efficiency of site 1 was significantly differential after editing in four varieties and the efficiency in Shaw was the highest (Fig. 4d). At the same time, the expression level of TraesCS3A02G263900 was down-regulated in HC374, Nyubai and Shaw, of which the expression level was significantly differential in Shaw after editing (Fig. 4e). These results suggest that the change of editing gene expression level may be caused by the change of miRNA binding ability caused by editing site. In general, the stronger the binding ability of miRNA, the weaker the gene expression. Meanwhile, there may be other regulatory mechanisms for the change of the expression level of this gene in Wuhan 1. Furthermore, the analysis of orthologues and conserved domain showed TraesCS3A02G263900 have a WD 40 domain and was also a orthologues gene of *OsRACK1* (Table S2 and S8). Component of the *OsRACK1* regulatory proteins that functions in innate immunity by interacting with multiple proteins in the RAC1 immune complex. *OsRACK1* also acts as positive regulator of reactive oxygen species (ROS) production and is required for resistance against rice blast (*M.grisea*) infection, indicating the potential function of TraesCS3A02G263900 in response to FHB.

**Fig. 4.**
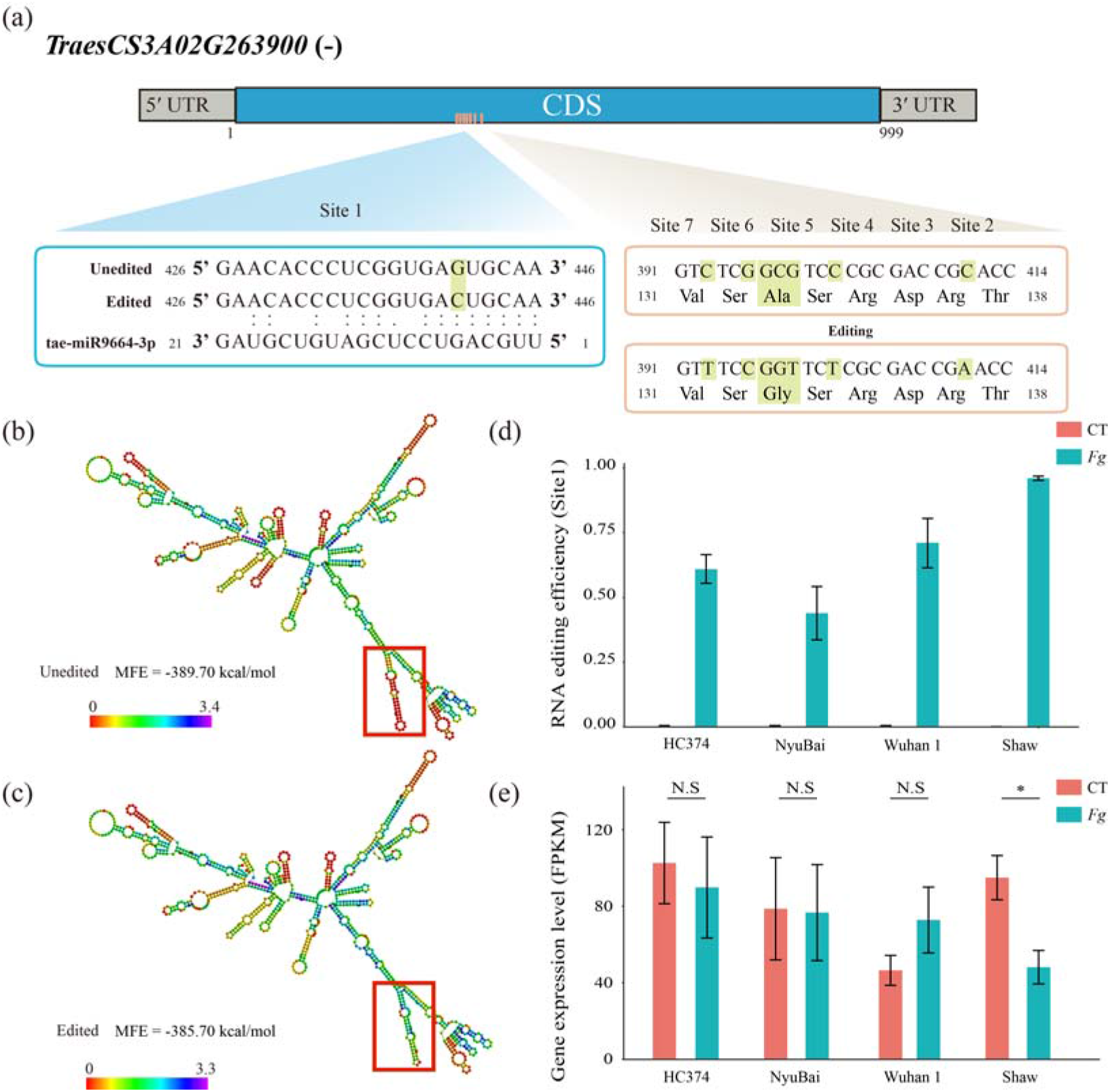
RNA editing effect on the miRNA targeting and RNA 2D structure on TraesCS3A02G263900. (a) Seven editing sites were identified in the coding region of TraesCS3A02G263900, of which site 4 and 5 was in the same amino acid and changed it, site 1 was found to changing the miRNA binding sites. (b) RNA secondary structure of TraesCS3A02G263900 before Site 1 editing. (c) RNA secondary structure of TraesCS3A02G263900 after Site 1 editing. (d) Editing efficiency of TraesCS1B02G294300 in four genotypes. (e) Expression levels of TraesCS1B02G294300 in four genotypes. *, P value < 0.05; **, P value < 0.01; ***, P value < 0.001; N.S, not significant

Tubulin is closely related to intracellular material transport, cell differentiation, cell movement, signal recognition, cell division and development. At the same time, plant tubulin is also related to the synthesis of cellulose microfibrils and plays a role in the growth and development of plant secondary wall (Yoshikawa et al. 2003). Here, TraesCS1D02G258800, belonging to Tubulin/FtsZ family and containing GTPase conserved domain (Table S8), were found to have nine RNA editing sites (1: chr1D_351247689; 2: chr1D_351247692; 3: chr1D_351247703; 4: chr1D_351247708; 5: chr1D_351247769; 6: chr1D_351247770; 7: chr1D_351247778; 8: chr1D_351247793; 9: chr1D_351247799) (Fig. 5a and Fig. S5). Among these sites, site 1, 2, 3, 4 and 6 changed the amino acid from Ser to Ala, Ile to Val, Met to Ile, Arg to Lys and Gly to Ser, respectively (Table S2). Furthermore, the RNA secondary and protein 3D structure prediction showed differences after editing. Due editing, the MFE of RNA secondary structure changed from −515.30 to −507.10 kcal/mol, indicating the stability of RNA structure decreased (Fig. 5b and Fig. 5c). The torsion of protein 3D structure changed from −1.81 to −2.20 (Fig. 5d and Fig. 5e).

**Fig. 5.**
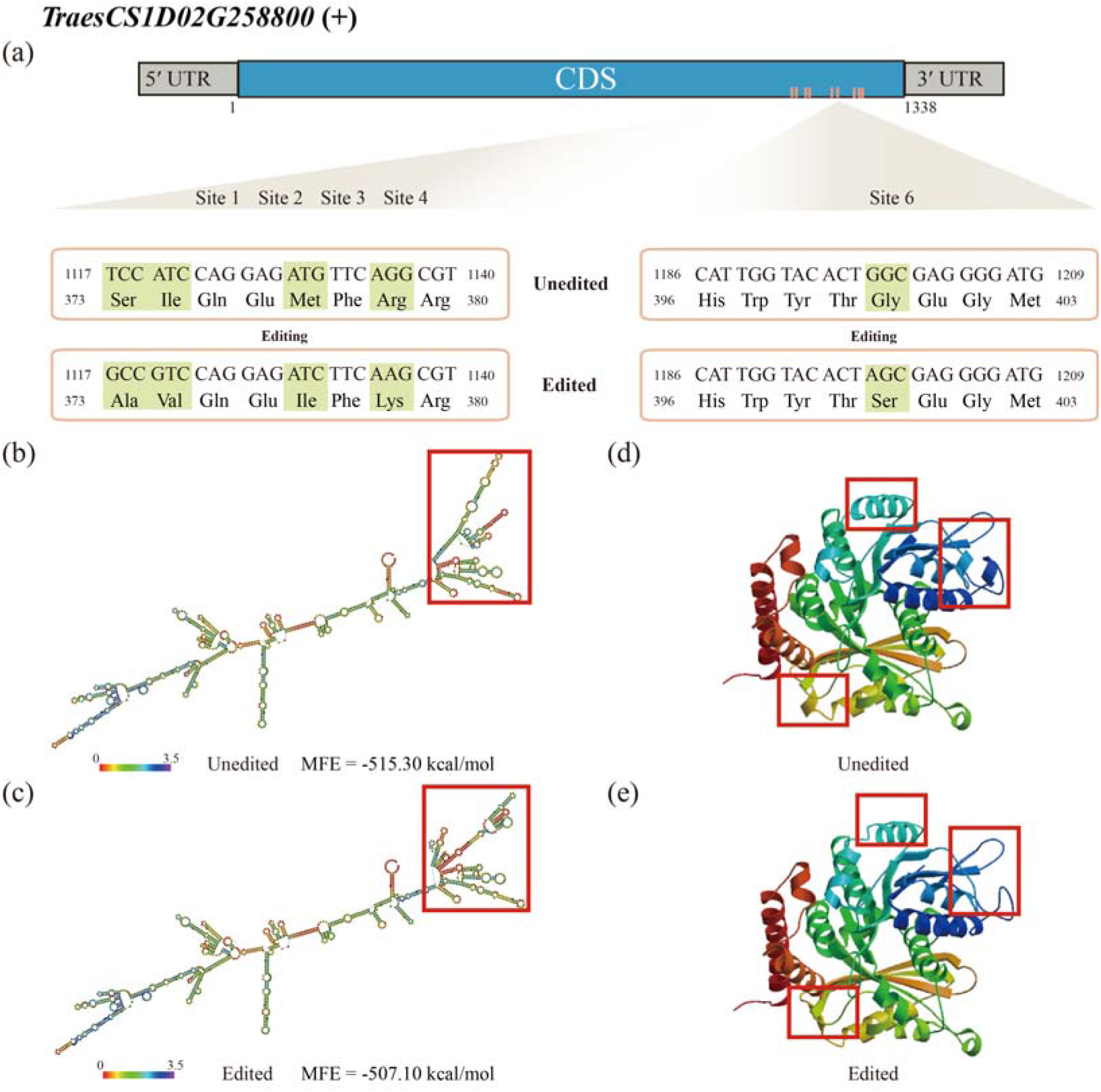
RNA editing effect on the mRNA 2D structure and protein 3D structure on TraesCS1D02G258800. (a) There were nine RNA editing sites found in the coding region of TraesCS1D02G258800, of which site 1, 2, 3, 4 and 6 caused the amino acid change and then also changed its 3D structure. (b-c) RNA secondary structure of TraesCS1D02G258800 before and after editing. (d-e) Protein 3D structure of TraesCS1D02G258800 before and after editing

## Conclusion

This is the first study to identify DNA/RNA differences associated with *Fg* infection in wheat at the whole transcriptome level. Totally, 187 unique DNA/RNA difference events (RNA editing sites) in 36 genes were identified in four varieties. The canonical G to A and C to T editing sites were found to be the most abundant, as well as other editing types were also identified. Integration of the RNA editing and gene expression, the differential expressed edited genes were also obtained, which could be considered as the potential resource for discovering the key novel genes associated *Fg* infection and tolerance. Finally, the effects of RNA editing were investigated and found that it could change the RNA secondary structure, protein 3D structure as well as miRNA targeting sites of edited genes to participate in the regulatory network of FHB response and tolerance. This study lay the foundation for further functional studies to reveal the roles of RNA editing playing in FHB response and tolerance in wheat, which will enrich the molecular basis underlying FHB tolerance, and also facilitate FHB tolerance improvement through epigenetic method in wheat and beyond.

## Abbreviations

*Fg*: Fusarium graminearum
FHB: Fusarium head blight
WGCNA: Weighted gene co-expression network analysis
dpi: Days post inoculation
DEG: Differentially expressed gene
DEEG: Differential expressed edited gene
GAPDH: Glyceraldehyde-3-phosphate dehydrogenase
DON: Deoxynivalenol
MFE: Minimum free energy
TSS: Translational start site
GO: Gene ontology
KEGG: Kyoto encyclopedia of genes and genomes

## Acknowledgement

We are grateful to Dr. Pan Youlian and his collaborators for sharing the RNA-seq data, which is publicly available from NCBI SRA database (SRP139946). We also appreciate the High-Performance Computing center of Northwest A&F University for providing computational resources.

## Data availability

The data that supports the findings of this study are available in the supplementary material of this article.

## Supplementary materials

**Fig. S1 Sample dendrogram and trait heatmap of WGCNA.** WGCNA was analyzed based on the expression level of the 58,380 expressed genes

**Fig. S2 Correlation of gene modules in WGCNA**

**Fig. S3 Correlation between gene modules and traits in WGCNA**

**Fig. S4 IGV results of RNA editing events in TraesCS3A02G263900.** (a) Site 1 in 4dpi of HC374. (b) Site 2-7 in 4dpi of HC374. (c) Site 1 in 4dpi of Nyubai. (d) Site 2-7 in 4dpi of Nyubai. (e) Site 1 in 4dpi of Wuhan 1. (f) Site 2-7 in 4dpi of Wuhan 1. (g) Site 1 in 4dpi of Shaw. (h) Site 2-7 in 4dpi of Shaw. The numbers on reads represent the editing efficiency of RNA editing sites in each sample. CT: Control group. Fg: Treatment group

**Fig. S5 IGV results of RNA editing events in TraesCS1D02G258800.** (a) Site 1-4 in 4dpi of HC374. (b) Site 5-9 in 4dpi of HC374. (c) Site 1-4 in 4dpi of Nyubai. (d) Site 5-9 in 4dpi of Nyubai. (e) Site 1-4 in 4dpi of Wuhan 1. (f) Site 5-9 in 4dpi of Wuhan 1. (g) Site 1-4 in 4dpi of Shaw. (h) Site 5-9 in 4dpi of Shaw. The numbers on reads represent the editing efficiency of RNA editing sites in each sample. CT: Control group. Fg: Treatment group

**Table S1. Summary of the identified DNA/RNA difference sites in this study.**

**Table S2. Message of RNA editing sites.**

**Table S3. Differential expressed level of RNA editing genes.**

**Table S4. Distribution of RNA editing genes in co-expression modules.**

**Table S5. Function enrichment of RNA editing genes.**

**Table S6. Minimum free energy (MFE) of RNA secondary structure of candidate editing genes.**

**Table S7. Prediction of the miRNA targeting sites of the RNA editing genes.**

**Table S8. Prediction of the conserved domain of the RNA editing genes.**

**Fig. S1.**
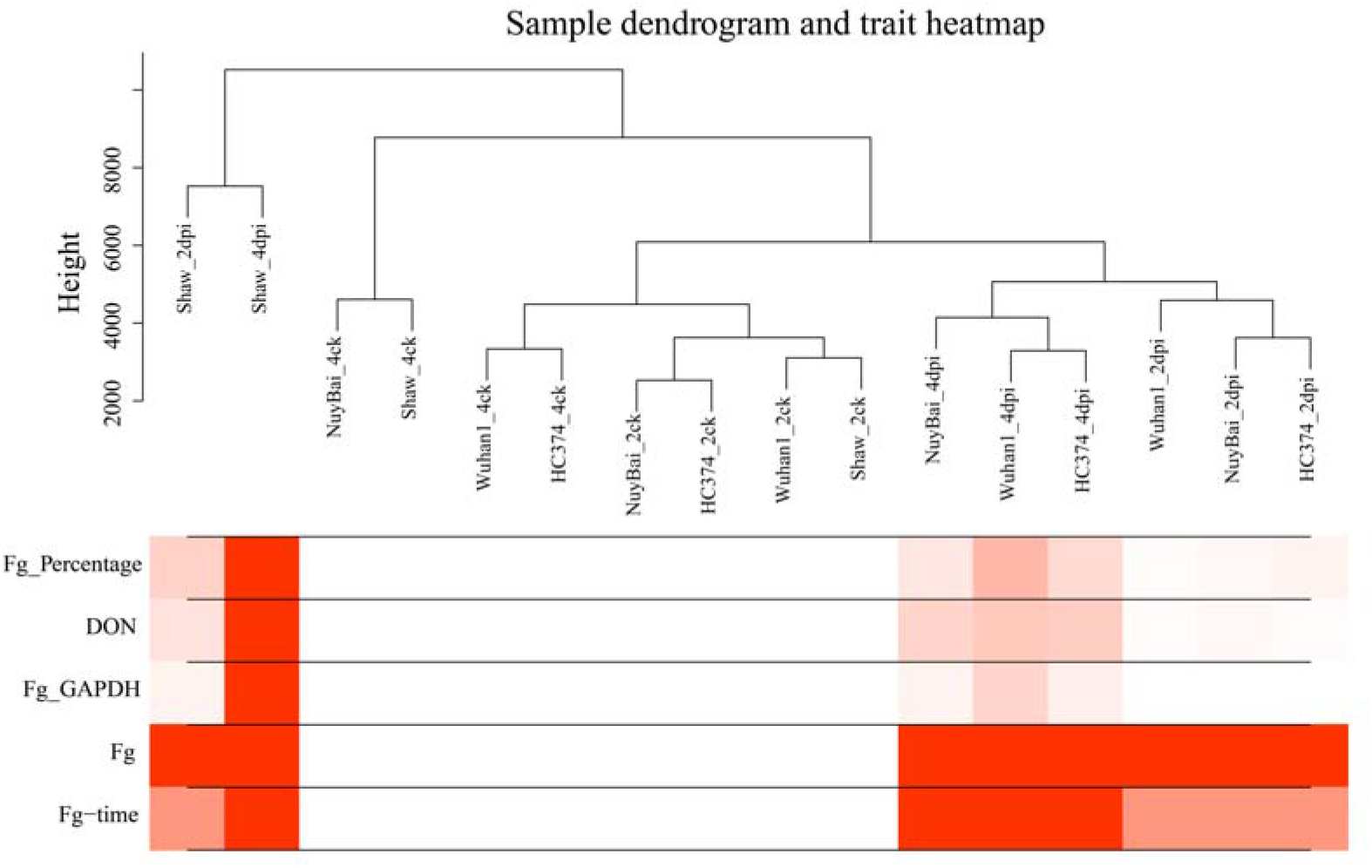
Sample dendrogram and trait heatmap of WGCNA. WGCNA was analyzed based on the expression level of the 58,380 expressed genes

**Fig. S2.**
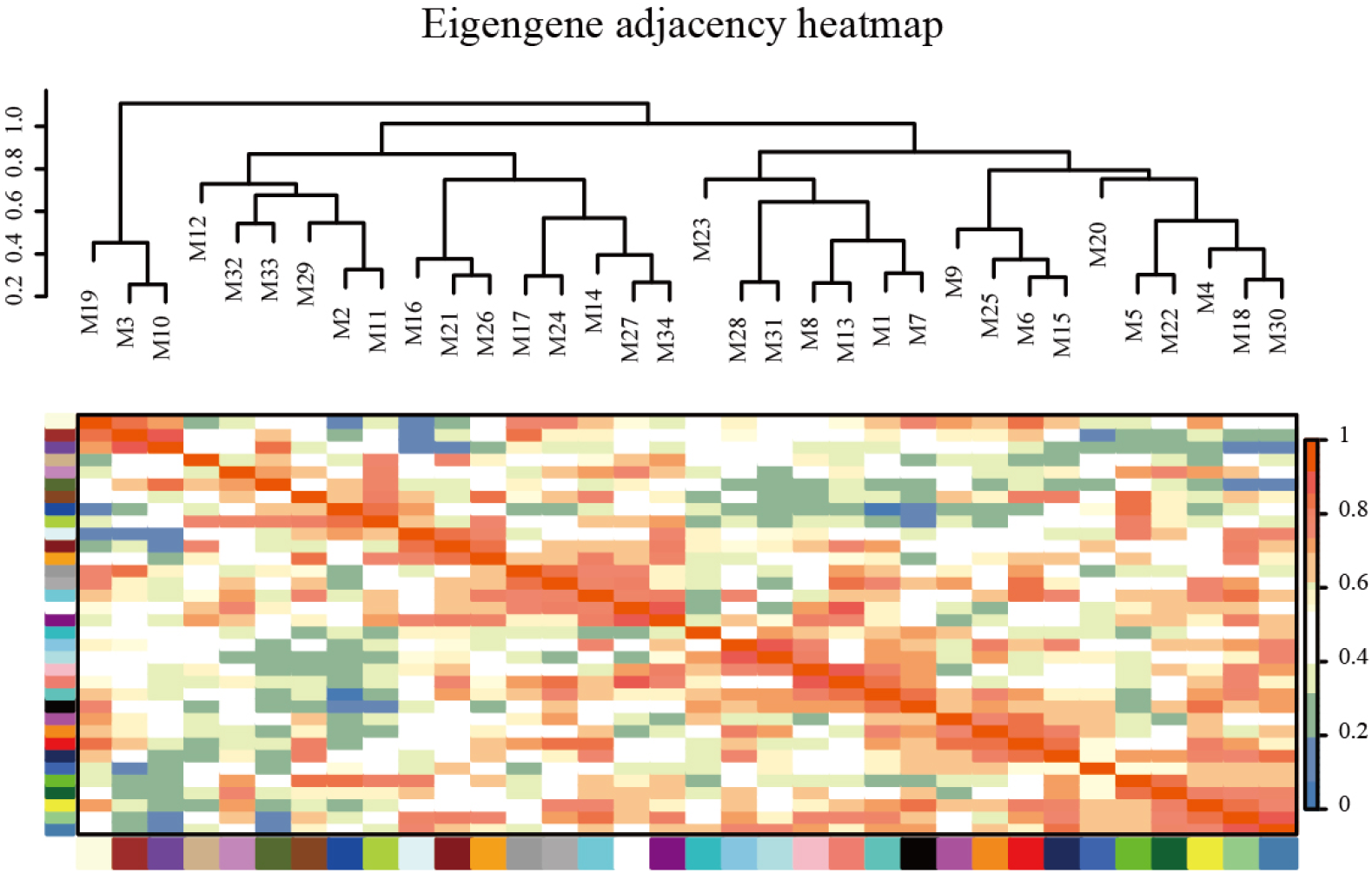
Correlation of gene modules in WGCNA.

**Fig. S3.**
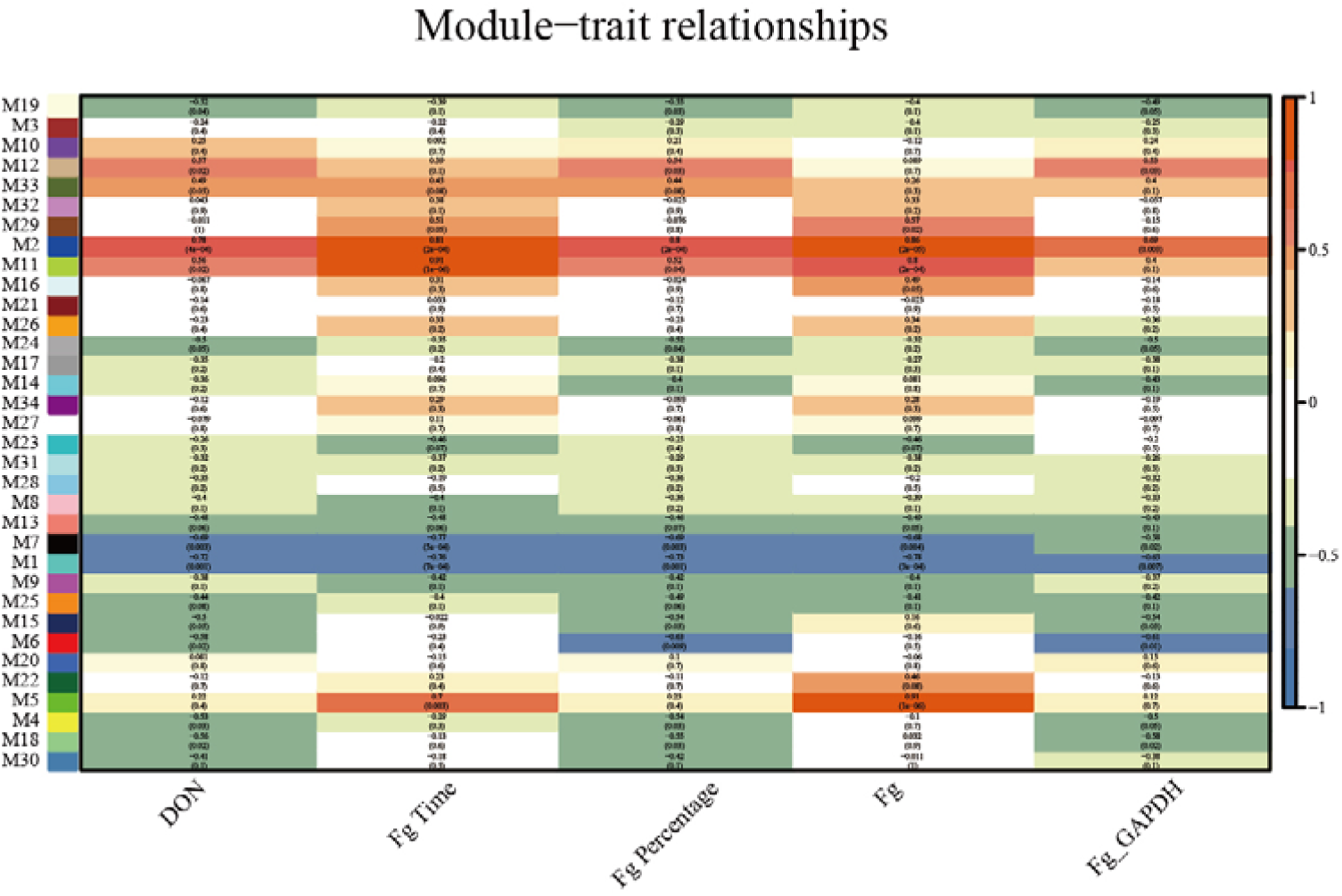
Correlation between gene modules and traits in WGCNA.

**Fig. S4.**
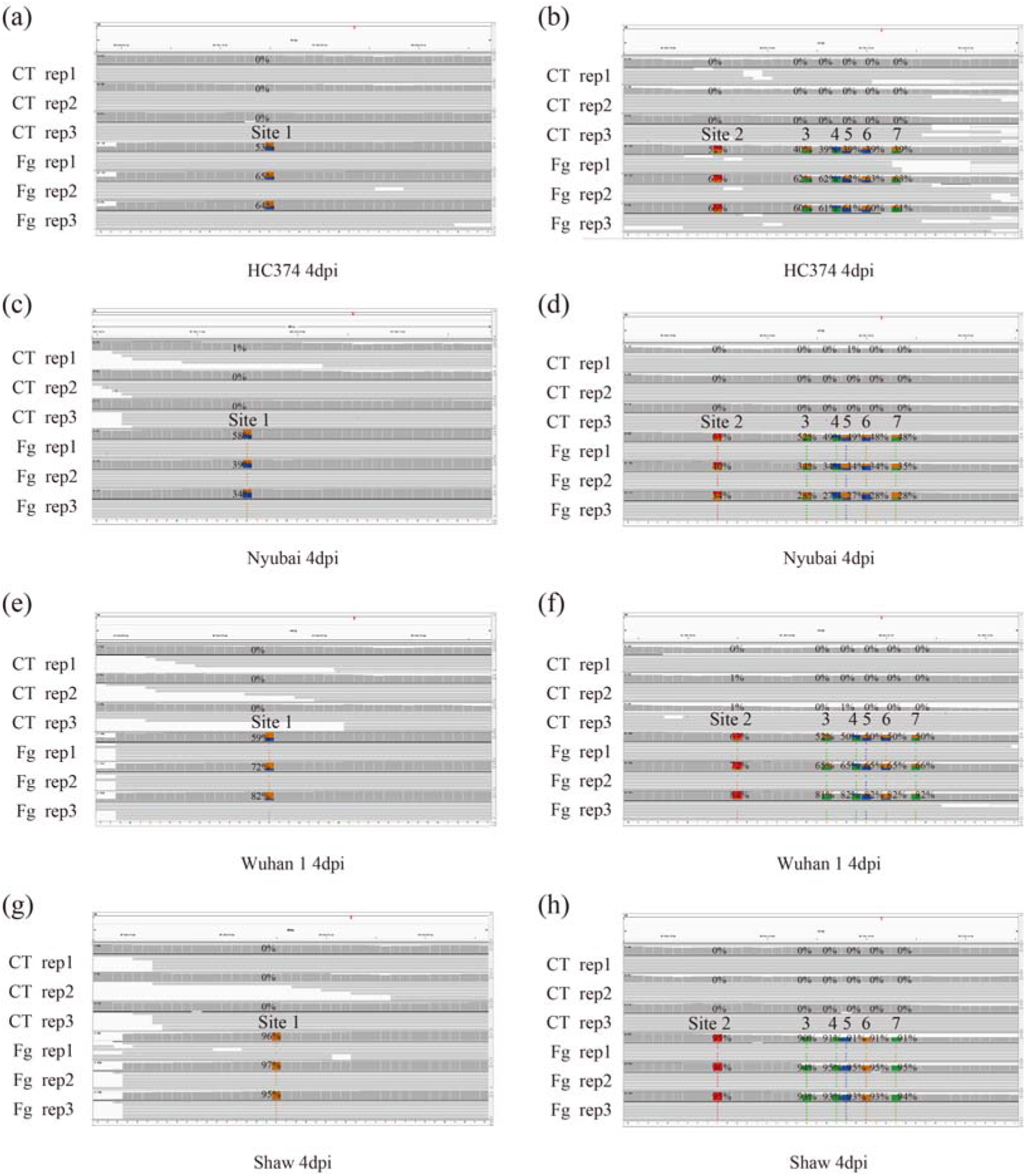
IGV results of RNA editing events in TraesCS3A02G263900. (a) Site 1 in 4dpi of HC374. (b) Site 2-7 in 4dpi of HC374. (c) Site 1 in 4dpi of Nyubai. (d) Site 2-7 in 4dpi of Nyubai. (e) Site 1 in 4dpi of Wuhan 1. (f) Site 2-7 in 4dpi of Wuhan 1. (g) Site 1 in 4dpi of Shaw. (h) Site 2-7 in 4dpi of Shaw. The numbers on reads represent the editing efficiency of RNA editing sites in each sample. CT: Control group. Fg: Treatment group

**Fig. S5.**
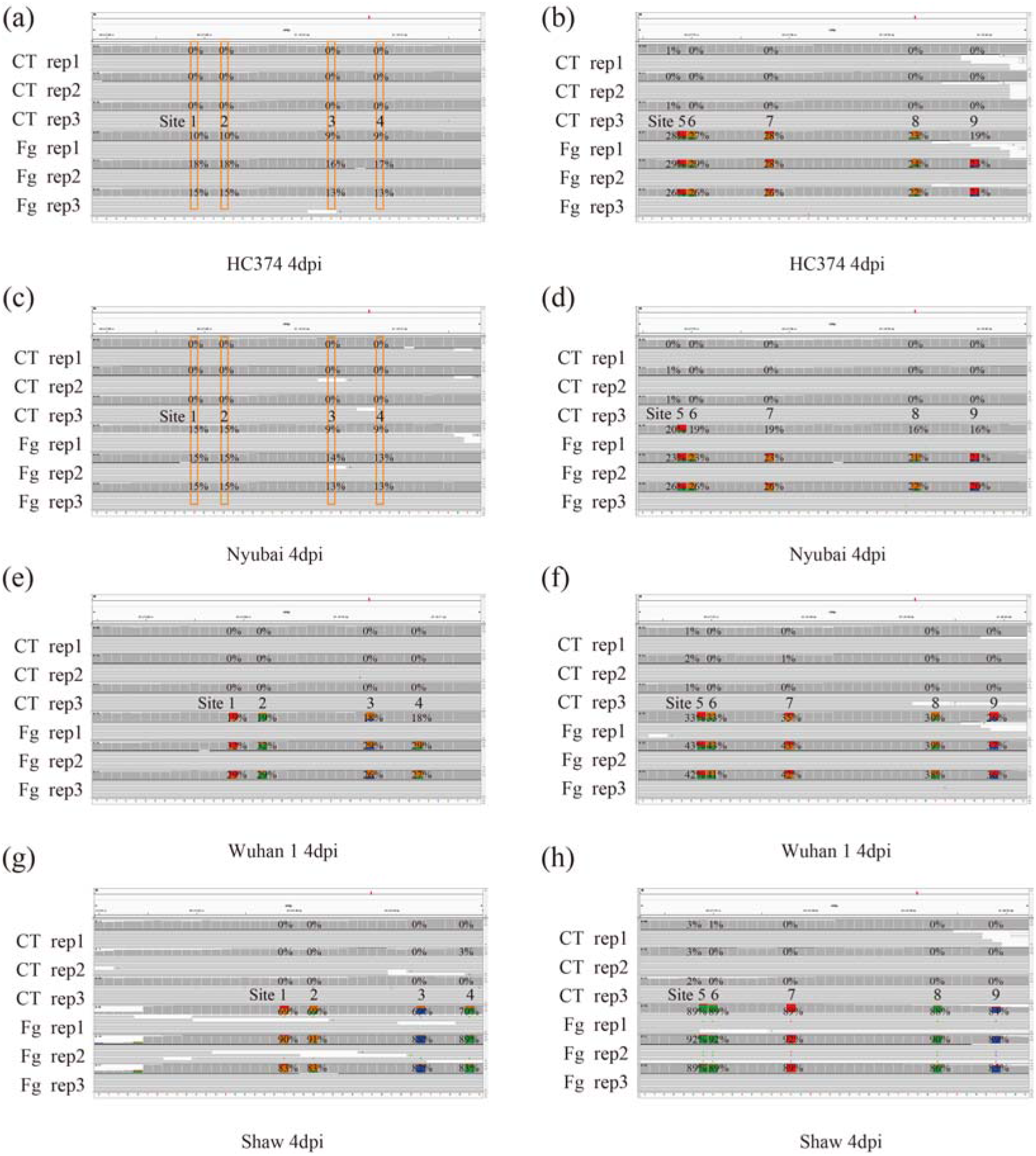
IGV results of RNA editing events in TraesCS1D02G258800. (a) Site 1-4 in 4dpi of HC374. (b) Site 5-9 in 4dpi of HC374. (c) Site 1-4 in 4dpi of Nyubai. (d) Site 5-9 in 4dpi of Nyubai. (e) Site 1-4 in 4dpi of Wuhan 1. (f) Site 5-9 in 4dpi of Wuhan 1. (g) Site 1-4 in 4dpi of Shaw. (h) Site 5-9 in 4dpi of Shaw. The numbers on reads represent the editing efficiency of RNA editing sites in each sample. CT: Control group. Fg: Treatment group

